# Intelligible speech synthesis from neural decoding of spoken sentences

**DOI:** 10.1101/481267

**Authors:** Gopala K. Anumanchipalli, Josh Chartier, Edward F. Chang

## Abstract

The ability to read out, or decode, mental content from brain activity has significant practical and scientific implications^1^. For example, technology that translates cortical activity into speech would be transformative for people unable to communicate as a result of neurological impairment^2,3,4^. Decoding speech from neural activity is challenging because speaking requires extremely precise and dynamic control of multiple vocal tract articulators on the order of milliseconds. Here, we designed a neural decoder that explicitly leverages the continuous kinematic and sound representations encoded in cortical activity^5,6^ to generate fluent and intelligible speech. A recurrent neural network first decoded vocal tract physiological signals from direct cortical recordings, and then transformed them to acoustic speech output. Robust decoding performance was achieved with as little as 25 minutes of training data. Naïve listeners were able to accurately identify these decoded sentences. Additionally, speech decoding was not only effective for audibly produced speech, but also when participants silently mimed speech. These results advance the development of speech neuroprosthetic technology to restore spoken communication in patients with disabling neurological disorders.

## Text

Neurological conditions that result in the loss of communication are devastating. Many patients rely on alternative communication devices that measure residual nonverbal movements of the head or eyes^7^, or even direct brain activity^8,9^, to control a cursor to select letters one-by-one to spell out words. While these systems dramatically enhance a patient’s quality of life, most users struggle to transmit more than 10 words/minute^10^, a rate far slower than the average of 150 words/min in natural speech. A major hurdle is how to overcome the constraints of current spelling-based approaches to enable far higher communication rates.

A promising alternative to spelling-based approaches is to directly synthesize speech^11,12^. Spelling is a sequential concatenation of discrete letters, whereas speech is produced from a fluid stream of overlapping, multi-articulator vocal tract movements^13^. For this reason, a biomimetic approach that focuses on vocal tract movements and the sounds they produce may be the only means to achieve the high communication rates of natural speech, and likely the most intuitive for users to learn^14,15^. In patients with paralysis, for example from ALS or brainstem stroke, high fidelity speech control signals may only be accessed by directly recording from intact cortical networks using a brain-computer interface.

Our goal was to demonstrate the feasibility of a neural speech prosthetic by translating brain signals into intelligible synthesized speech at the rate of a fluent speaker. To accomplish this, we recorded high-density electrocorticography (ECoG) signals from three participants undergoing intracranial monitoring for epilepsy treatment as they spoke several hundred sentences aloud. We designed a recurrent neural network that decoded cortical signals with an explicit intermediate representation of the articulatory dynamics to generate audible speech.

An overview of our two-stage decoder approach is shown in Figure 1a-d. In the first stage, a bidirectional long short term memory (bLSTM) recurrent neural network^16^ decodes articulatory kinematic features from continuous neural activity (Figure 1a, b). In the second stage, a separate bLSTM decodes acoustic features from the decoded articulatory features from stage 1 (Figure 1c). The audio signal is then synthesized from the decoded acoustic features (Figure 1d).

**Figure 1:**
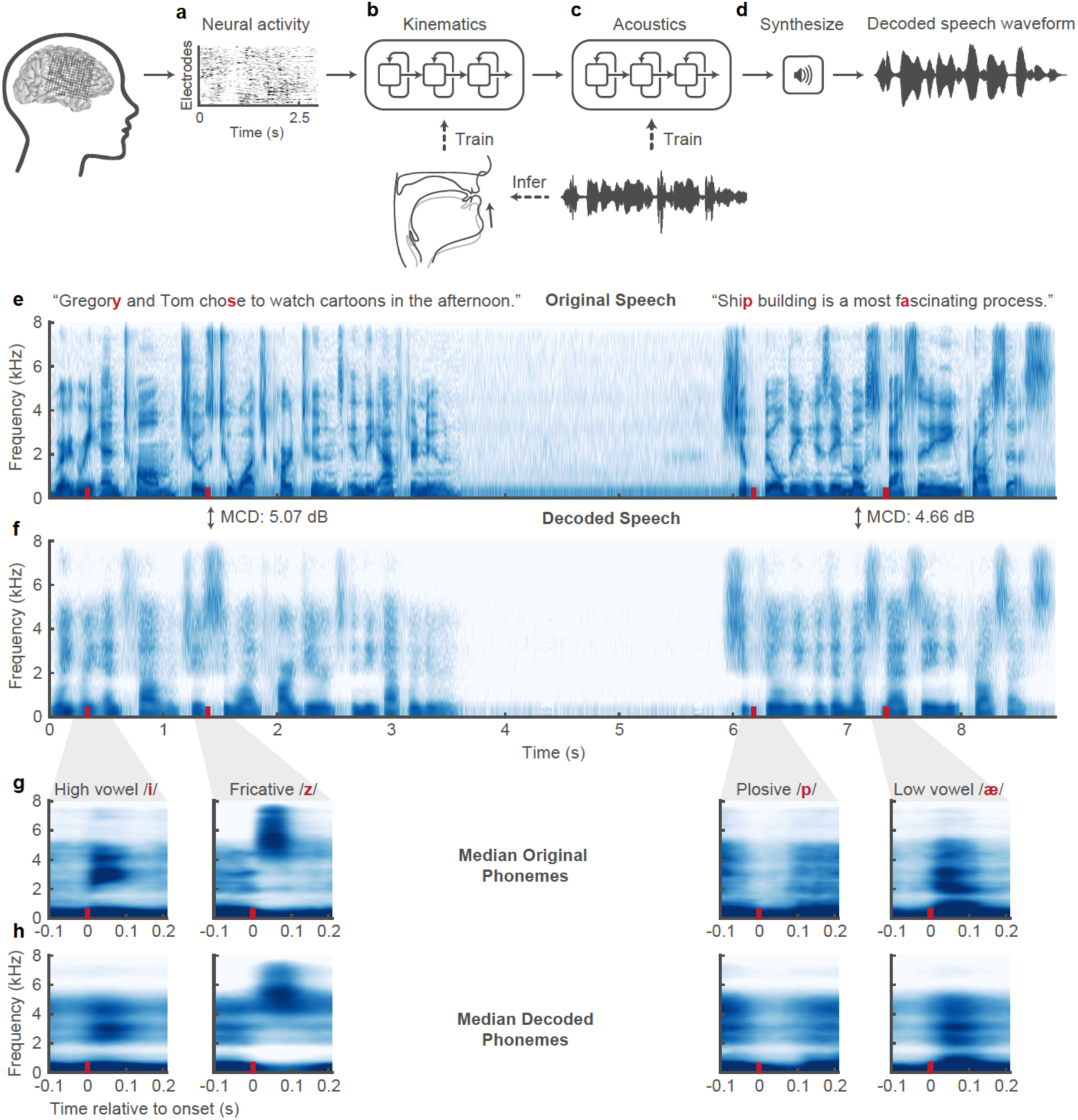
Speech synthesis from neurally decoded spoken sentences. **a**, The neural decoding process begins by extracting high-gamma amplitude (70-200Hz) and low- frequency (1-30Hz) ECoG activity. **b**, A 3-layer bi-directional long short term memory (bLSTM) neural network learns to decode kinematic representations of articulation from filtered ECoG signals. **c**, An additional 3-layer bLSTM learns to decode acoustics from the previously decoded kinematics. Acoustics are represented as spectral features (e.g. Mel-frequency cepstral coefficients (MFCCs)) extracted from the speech waveform. **d**, Decoded signals are synthesized into an acoustic waveform. **e**, Spectrogram shows the frequency content of two sentences spoken by a participant. **f**, Spectrogram of synthesized speech from brain signals recorded simultaneously with the speech in **e**. Mel- cepstral distortion (MCD), a metric for assessing the spectral distortion between two audio signals, was computed for each sentence between the original and decoded audio. **g,h** 300 ms long, median spectrograms that were time-locked to the acoustic onset of phonemes from original (**g**) and decoded (**h**) audio. Medians were computed from phonemes in 100 sentences that were withheld during decoder training (n: /i/ = 112, /z/ = 115, /p/ 69, /ae/ = 86). These phonemes represent the diversity of spectral features. Original and decoded median phoneme spectrograms were well correlated (r > 0.9 for all phonemes, p=1e-18)

There are three sources of data for training the decoder: high density ECoG recordings, acoustics, and articulatory kinematics. For ECoG, high-gamma amplitude envelope (70-200 Hz)^17^, and low frequency component (1-30 Hz)^18^ were extracted from the raw signal of each electrode. Electrodes were selected if they were located on key cortical areas for speech: ventral sensorimotor cortex (vSMC)^19^, superior temporal gyrus (STG)^20^, or inferior frontal gyrus (IFG)^21^ (Figure 1a). For acoustics, instead of a typical spectrogram, we used 25 mel-frequency cepstral coefficients (MFCCs), 5 sub-band voicing strengths for glottal excitation modelling, pitch, and voicing (32 features in all). These acoustic parameters are specifically designed to emphasize perceptually relevant acoustic features while maximizing audio reconstruction quality^22^.

Lastly, a key component of our decoder is an intermediate articulatory kinematic representation between neural activity and acoustics (Figure 1b). Our previous work demonstrated that articulatory kinematics is the predominant representation in the vSMC^6^. Since it was not possible to record articulatory movements synchronously with neural recordings, we used a statistical speaker-independent Acoustic-to-Articulatory inversion method to estimate vocal tract kinematic trajectories corresponding to the participant’s produced speech acoustics. We added additional physiological features (e.g. manner of articulation) to complement the kinematics and optimized these values within a speech autoencoder to infer the full intermediate articulatory kinematic representation that captures vocal tract physiology during speech production (see methods). From these features, it was possible to accurately reconstruct the speech spectrogram (Figure 1e,f).

### Synthesis performance

Overall, we observed highly detailed reconstructions of speech decoded from neural activity alone (See supplemental video). Examples of decoding performance are shown in Figure 1 (e,f), where the audio spectrograms from two original spoken sentences are plotted above those decoded from brain activity. The first sentence is representative of the median performance and the second shows one of the best decoded sentences. The decoded spectrogram contained salient energy patterns present in the original spectrogram.

To illustrate the quality of reconstruction at the phonetic level, we compared median spectrograms of phonemes from original and decoded audio. As shown in Figure 1 g,h, the formant frequencies (F1-F3, seen as high energy resonant bands in the spectrograms) and distribution of spectral energy for high and low vowels (/i/ and /ae/, respectively) of the decoded examples closely resembled the original speech. For alveolar fricatives (/z/) the high frequency (>4kHz) acoustic energy was well represented in both spectrograms. For plosives (/p/), the short pause (relative silence during the closure) followed by a broadband burst of energy (after the release) was also well decoded. The decoder also correctly reconstructed the silence in between the sentences when the participant was not speaking.

To quantify performance, we tested the neural decoder for each participant on 100 sentences that were withheld during the training and optimization of the full model. In traditional speech synthesis, the spectral distortion of synthesized speech from ground-truth is commonly reported using the mean Mel-Cepstral Distortion (MCD)^23^. The use of Mel-Frequency bands emphasizes the distortion of perceptually relevant frequency bands of the audio spectrogram^24^. In Figure 2a, the MCD of neurally decoded speech was compared with reference synthesis from articulatory kinematics and chance-level decoding (lower MCD is better). The reference synthesis acts as a bound for performance as it simulated what perfect neural decoding of the kinematics would achieve. For our participants (P1, P2, P3), the median MCD scores of decoding speech were 5.14 dB, 5.55 dB, and 5.49 dB, all better than chance-level decoding (p<1e-18, n=100 sentences, Wilcoxon signed-rank test (WSRT), for each participant). These scores were on par with state-of-the-art approaches to decode speech from facial surface electromyography (EMG) with similarly sized datasets (average MCD of 5.21 dB)^25^.

**Figure 2:**
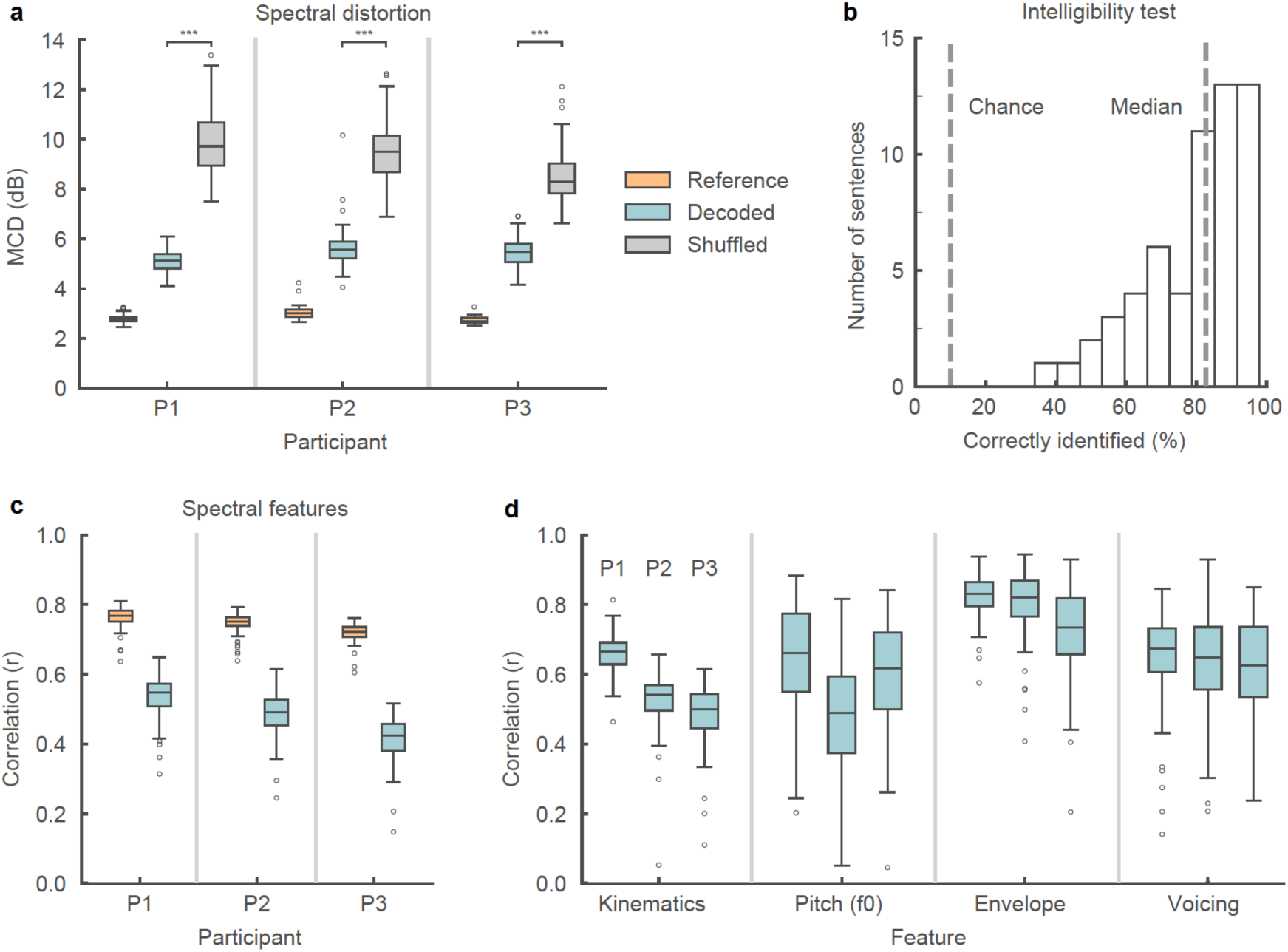
Decoded speech intelligibility and feature-specific performance.. **a**, Spectral distortion, measured by Mel-Cepstral Distortion (MCD) (lower values are better), between original spoken sentences and neurally decoded sentences that were held out from model training (n = 100). Reference MCD refers to the MCD resulting from the synthesis of original kinematics without neural decoding and provides an upper bound for performance. MCD scores were compared to chance-level MCD scores obtained by shuffling data before decoding. **b**, Decoded sentence intelligibility was assessed by asking naïve participants to identify the sentence they heard from 10 choices. Each sample (n = 60) represents the percentage of correctly identified trials for one sentence. The median sentence was correctly identified 83% of the time. **c**, Correlation of original and decoded spectral features. Values represent the mean correlation of the 32 spectral features for each sentence (n = 100). Correlation performance for individual spectral features is reported in extended data figure 1b. **d**, Correlations between original and decoded intelligibility-relevant features. Kinematic values represent the mean correlation of the 33 kinematic features (the intermediate representation) for each sentence (n =100). Correlation performance for individual kinematic features is reported in extended data figure 1a. Box plots depict median (horizontal line inside box), 25th and 75th percentiles (box), 25/75th percentiles ±1.5× interquartile range (whiskers), and outliers (circles). Distributions were compared with each as other as indicated or with chance-level distributions using two-tailed Wilcoxon signed-rank tests (p < 1e-10, n = 100, for all tests).

To assess the perceptual intelligibility of the decoded speech, we used Amazon Mechanical Turk to evaluate naïve listeners’ ability to understand the neurally decoded trials. We asked 166 people to identify which of 10 sentences (written on screen) corresponded to the decoded audio they heard. The median percentage of participants who correctly identified each sentence was 83%, significantly above chance (10%) (Figure 2b).

In addition to spectral distortion and intelligibility, we also examined the correlations between original and decoded spectral features. The median correlations (of sentences, Pearson’s r) of the mean decoded spectral feature (pitch + 25 MFCCs + excitation strengths + voicing) for each participant were 0.55, 0.49, and 0.42 (Figure 2c). Similarly, for decoded kinematics (the intermediate representation), the median correlations were 0.66, 0.54, and 0.50 (Figure 2d). Finally, we examined three key aspects of prosody for intelligible speech: pitch (f0), speech envelope, and voicing^26^ (Figure 2d). For all participants, these features were decoded well above chance-level correlations (r > 0.6, except f0 for P2: r= 0.49, p<1e-10, n=100, WSRT, for all participants and features in Figure 2c-d). Correlation decoding performance for all other features is shown in Extended Data Figure 1a,b.

### Effects of model design decisions

The following analyses were performed on data from P1. In designing a neural decoder for clinical applications, there are several key considerations regarding the input to the model. First, in patients with severe paralysis or limited speech ability, training data may be very difficult to obtain. In audio-based commercial applications like digital assistants, successful speech synthesis from text relies on tens of hours of speech^27^. Despite having limited neural data, we observed high decoding performance, and therefore we wanted to assess how much data was necessary to achieve this level of performance. Furthermore, we wanted to see if there was a clear advantage in explicitly modeling articulatory kinematics as an intermediate step over decoding acoustics directly from the ECoG signals. The motivation for including articulatory kinematics was to reduce the complexity of the ECoG-to-acoustic mapping because it captures the physiological process by which speech is generated and is encoded in the vSMC^6^.

We found robust performance could be achieved with as little as 25 minutes of speech, but performance continued to improve with the addition of more data (Figure 3a,b). A crucial factor in performance was the articulatory intermediate training step. Without this step, direct ECoG to acoustic decoding MCD was offset by 0.54 dB using the full data set (Figure 3a) (p=1e-17, n=100, WSRT), a substantial difference given that a change in MCD as small as 0.2 dB is perceptually noticeable^28^. While the two approaches might perform comparably with enough data, the biomimetic approach using an intermediate articulatory representation is superior because it requires less training data.

**Figure 3:**
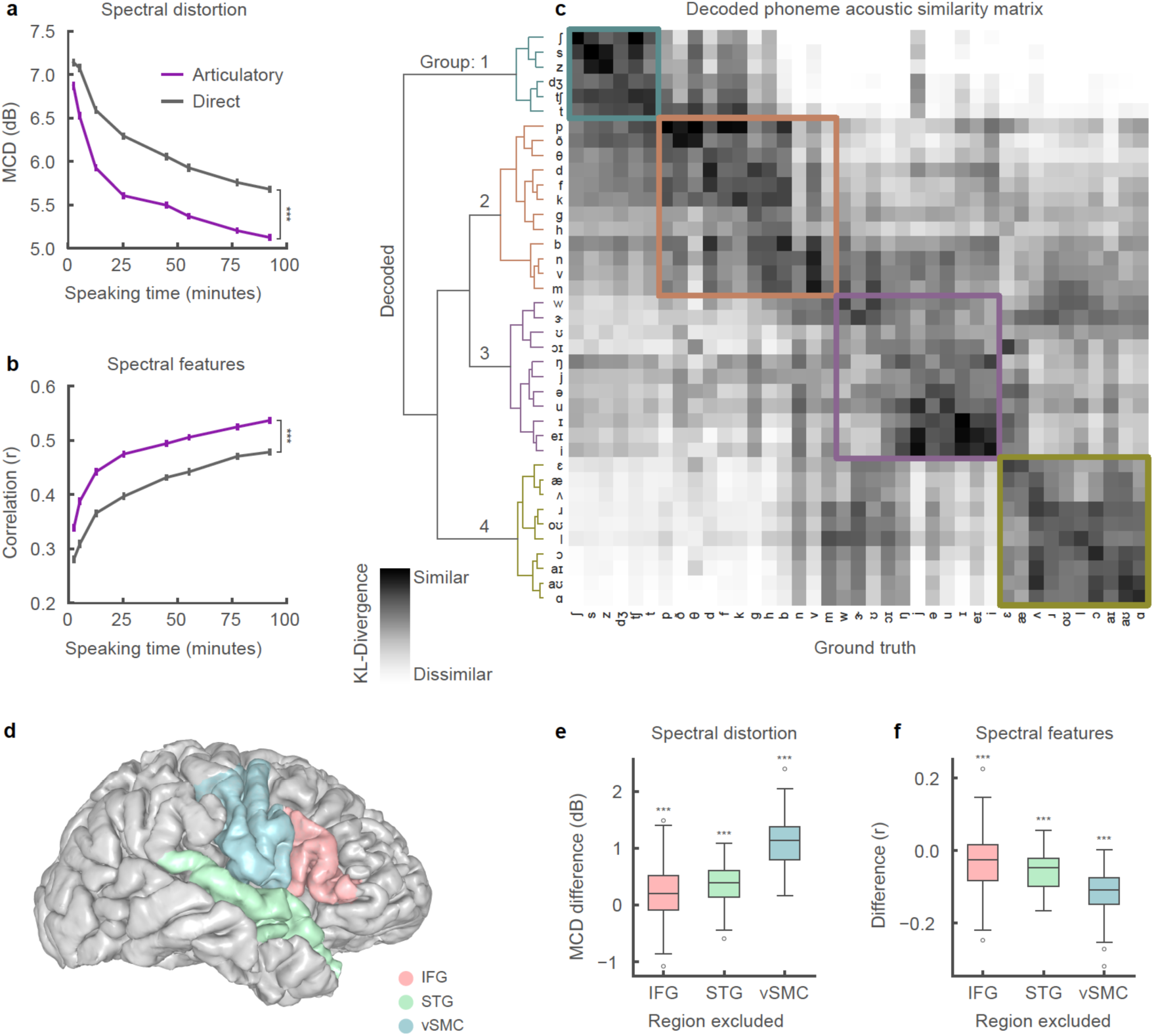
Effects of model design decisions. **a**, **b**, Mean correlation of original and decoded spectral features (**a)** and mean spectral distortion (MCD) (**b**) for model trained on varying amounts of training data. Training data was split according to recording session boundaries resulting the following sizes: 2.4, 5.2, 12.6, 25.3, 44.9, 55.2, 77.4, and 92.3 minutes of speaking data. The neural decoding approach that included an articulatory intermediate stage (purple) performed significantly better with every size of training data than direct ECoG to acoustics decoder (grey) (all: p < 1e-5, n = 100; Wilcoxon signed-rank test, error bars = SE). **c**, Acoustic similarity matrix compares acoustic properties of decoded phonemes and originally spoken phonemes. Similarity is computed by first estimating a gaussian kernel density for each phoneme (both decoded and original) and then computing the Kullback-Leibler (KL) divergence between a pair of decoded and original phoneme distributions. Each row compares the acoustic properties of a decoded phoneme with originally spoken phonemes (columns). Hierarchical clustering was performed on the resulting similarity matrix. **d**, Anatomical reconstruction of a single participant’s brain with the following regions used for neural decoding: ventral sensorimotor cortex (vSMC), superior temporal gyrus (STG), and inferior frontal gyrus (IFG). **e**, **f**, Difference in spectral distortion (MCD) (**e**), and difference in correlation (Pearson’s r) performance (**f**) between decoder trained on all regions and decoders trained on all-but-one region. Exclusion of any region resulted in decreased performance (p < 3e-4, n = 100; Wilcoxon signed-rank test). Box plots as described in Figure 2.

Second, we wanted to understand the acoustic-phonetic properties that were preserved in decoded speech because they are important for relative phonetic discrimination. To do this, we compared the acoustic properties of decoded phonemes to ground truth by constructing a statistical distribution of the spectral feature vectors for each phoneme. Using Kullback-Leibler (KL) divergence, we compared the distribution of each decoded phoneme to the distribution of each ground-truth phoneme to determine how similar they were (Figure 3c). From the acoustic similarity matrix of only ground-truth phoneme-pairs (Extended Data Figure 2), we expected that, in addition to the same decoded and ground-truth phoneme being similar to one another, phonemes with shared acoustic properties would also be characterized as similar to one another. For example, two fricatives will be more acoustically similar to one another than to a vowel.

Hierarchical clustering on the KL-divergence of each phoneme pair demonstrated that phonemes were clustered into four main groups. These groups represent the primary decoded acoustic differences between phonemes. Within each group, phonemes were more likely to be confused with one another due to their shared acoustic properties. For instance, a decoded /s/ may easily be confused with /z/ or other phonemes in Group 1. Group 1 contained consonants with an alveolar place of constriction. Group 2 contained almost all other consonants. Group 3 contained mostly high vowels. Group 4 contained mostly mid and low vowels. The difference between groups tended to correspond to variations along acoustically significant dimensions (frequency range of spectral energy for consonants, and formants for vowels). These groupings were similar to those obtained by clustering KL-divergence of ground-truth phoneme pairs (Extended Data Figure 2).

Third, since the success of the decoder depends on the initial electrode placement, we wanted to assess how much the cortical activity of each brain region contributed to decoder performance. We quantified the contributions of the vSMC, STG, and IFG by training decoders in a leave-one-region-out fashion and comparing performance (Figure 3d). Removing any region led to decreased decoder performance (Figure 3e-f) (p<3e-4, n=100, WSRT). However, excluding vSMC resulted in the largest decrease in performance (1.13 dB MCD increase).

### Silently mimed speech decoding

Finally, since future speech decoding applications must work even when speakers do not produce audible sounds, we tested our decoder with a held-out set of 58 sentences in which the participant (P1) audibly produced each sentence and then mimed the same sentence, making the same kinematic movements but without making sound. Even though the decoder was not trained on any mimed speech, the spectrograms of synthesized silent speech demonstrated similar spectral features when compared to synthesized audible speech of the same sentence (Figure 4a-c). After dynamic time warping the acoustics of the decoded silent speech with the original audio of the preceding audibly produced sentence, we calculated the spectral distortion and correlation of the spectral features (Figure 4d,e). As expected, performance on mimed speech was inferior to spoken speech (30% MCD difference) although this is consistent with earlier work on silent facial EMG-to-speech synthesis where decoding performance from EMG signals was significantly worse when participants silently articulated without audible speech output^29^. The performance gap may also be due to the absence of voicing and laryngeal activation. This demonstrates that it is possible to decode important spectral features of speech that were never audibly uttered (p < 1e-11, compared to chance, n = 58; Wilcoxon signed-rank test).

**Figure 4:**
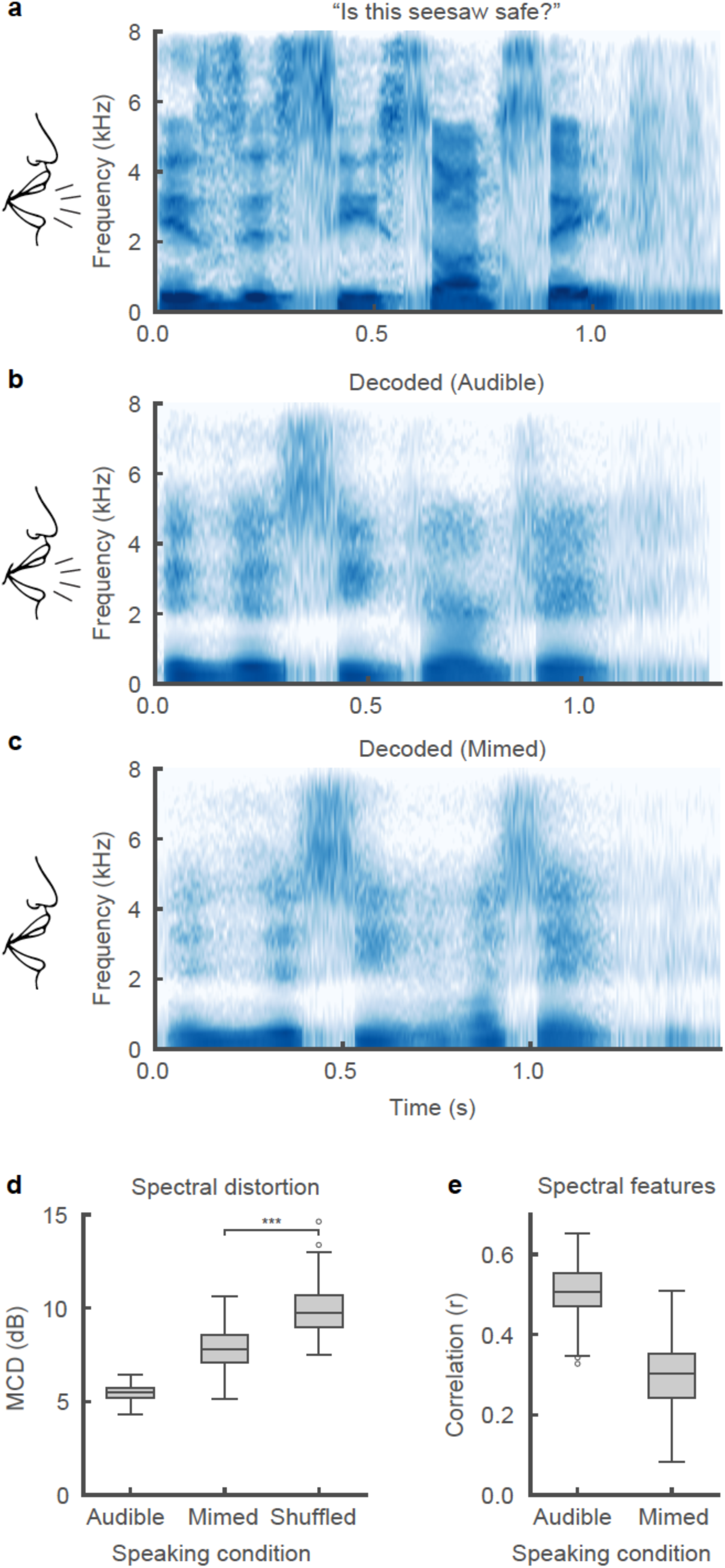
Speech synthesis from neural decoding of silently mimed speech. **a**-**c**, Spectrograms of original spoken sentence (**a**), neural decoding from audible production (**b**), and neural decoding from silently mimed production (**c**). **d**, **e**, Spectral distortion (MCD) (**d**) and correlation of original and decoded spectral features (**e**) for audibly and silently produced speech. Since correlations are with respect to original audibly produced sentences, decoded sentences that were silently mimed were dynamically time-warped according to their spectral features. Decoded sentences were significantly better than chance-level decoding for both speaking conditions (p < 1e-11, for all comparisons, n = 58; Wilcoxon signed-rank test). Box plots as described in Figure 2.

### Discussion

Our results demonstrate intelligible speech synthesis from ECoG during both audible and silently mimed speech production. Previous strategies for neural decoding of speech have primarily focused on direct classification of speech segments like phonemes or words^30,31,32,33^. However, these demonstrations have been limited in their ability to scale to larger vocabulary sizes and communication rates. Meanwhile, decoding of auditory cortex responses has been more successful for continuous speech sounds^18,34^, in part because of the direct relationship between the auditory encoding of spectrotemporal information and the reconstructed spectrogram. An outstanding question has been whether decoding vocal tract movements from the speech motor cortex could be used for generating high-fidelity acoustic output.

We believe that cortical activity at vSMC electrodes was critical for decoding (Figure 3e,f) because it encodes the underlying articulatory physiology that produces speech^6^. Our decoder explicitly incorporated this knowledge to simplify the complex mapping from neural activity to sound by first decoding the physiological correlate of neural activity and then transforming to speech acoustics. We have demonstrated that this statistical mapping permits generalization with limited amounts of training.

Direct speech synthesis has several major advantages over spelling-based approaches. In addition to the capability to communicate at a natural speaking rate, it captures prosodic elements of speech that are not available with text output, for example pitch intonation (Figure 2d) and word emphasis^35^. Furthermore, a practical limitation for current alternative communication devices is the cognitive effort required to learn and use them. For patients in whom the cortical processing of articulation is still intact, a speech-based BCI decoder may be far more intuitive and easier to learn to use^14,15^.

Brain-computer interfaces are rapidly becoming clinically viable means to restore lost function^36^. Impressive gains have already been made motor restoration of cursor control and limb movements. Neural prosthetic control was first demonstrated in participants without disabilities^37,38,39^ before translating the technology to participants with tetraplegia^40,41,42,43^. While this articulatory-based approach establishes a new foundation for speech decoding, we anticipate additional improvements from modeling higher-order linguistic and planning goals^44,45^. Our results may be an important next step in realizing speech restoration for patients with paralysis.

## Methods

### Participants and experimental task

Three human participants (30 F, 31 F, 34 M) underwent chronic implantation of high-density, subdural electrode array over the lateral surface of the brain as part of their clinical treatment of epilepsy (right, left, and right hemisphere grids, respectively). Participants gave their written informed consent before the day of the surgery. All participants were fluent in English. All protocols were approved by the Committee on Human Research at UCSF. Each participant read and/or freely spoke a variety of sentences. P1 read aloud two complete sets of 460 sentences from the MOCHA-TIMIT database^46^. Additionally, P1 also read aloud passages from the following stories: Sleeping Beauty, Frog Prince, Hare and the Tortoise, The Princess and the Pea, and Alice in Wonderland. P2 read aloud one full set of 460 sentences from the MOCHA-TIMIT database and further read a subset of 50 sentences an additional 9 times each. P3 read 596 sentences describing three picture scenes and then freely described the seen resulting in another 254 sentences. P3 also spoke 743 sentences during free response interviews. In addition to audible speech, P1 also read 10 sentences 12 times each alternating between audible and silent (mimed i.e. making the necessary mouth movements) speech. Microphone recordings were obtained synchronously with the ECoG recordings.

### Data acquisition and signal processing

Electrocorticography was recorded with a multi-channel amplifier optically connected to a digital signal processor (Tucker-Davis Technologies). Speech was amplified digitally and recorded with a microphone simultaneously with the cortical recordings. ECoG electrodes were arranged in a 16 × 16 grid with 4 mm pitch. The grid placements were decided upon purely by clinical considerations. ECoG signals were recorded at a sampling rate of 3,052 Hz. Each channel was visually and quantitatively inspected for artifacts or excessive noise (typically 60 Hz line noise). The analytic amplitude of the high-gamma frequency component of the local field potentials (70 - 200 Hz) was extracted with the Hilbert transform and down-sampled to 200 Hz. The low frequency component (1-30 Hz) was also extracted with a 5th order Butterworth bandpass filter and parallelly aligned with the high-gamma amplitude. Finally, the signals were z-scored relative to a 30 second window of running mean and standard deviation, so as to normalize the data across different recording sessions. We studied high-gamma amplitude because it has been shown to correlate well with multi-unit firing rates and has the temporal resolution to resolve fine articulatory movements^17^. We also included a low frequency signal component due to the decoding performance improvements note for reconstructing perceived speech from auditory cortex^34^. Decoding models were constructed using all electrodes from vSMC, STG, and IFG except for electrodes with bad signal quality as determined by visual inspection.

### Phonetic and phonological transcription

For the collected speech acoustic recordings, transcriptions were corrected manually at the word level so that the transcript reflected the vocalization that the participant actually produced. Given sentence level transcriptions and acoustic utterances chunked at the sentence level, hidden Markov model based acoustic models were built for each participant so as to perform sub-phonetic alignment^47^. Phonological context features were also generated from the phonetic labels, given their phonetic, syllabic and word contexts.

### Cortical surface extraction and electrode visualization

We localized electrodes on each individual’s brain by co-registering the preoperative T1 MRI with a postoperative CT scan containing the electrode locations, using a normalized mutual information routine in SPM12. Pial surface reconstructions were created using Freesurfer. Final anatomical labeling and plotting was performed using the img_pipe python package^48^.

### Inference of articulatory kinematics

The articulatory kinematics inference model comprises a stacked deep encoder-decoder, where the encoder combines phonological and acoustic representations into a latent articulatory representation that is then decoded to reconstruct the original acoustic signal. The latent representation is initialized with inferred articulatory movement from Electromagnetic Midsagittal Articulography (EMA)^6^ and appropriate manner features.

Chartier et al., 2018 described a statistical subject-independent approach to acoustic-to-articulatory inversion which estimates 12 dimensional articulatory kinematic trajectories (x and y displacements of tongue dorsum, tongue blade, tongue tip, jaw, upper lip and lower lip, as would be measured by EMA) using only the produced acoustics and phonetic transcriptions. Since, EMA features do not describe all acoustically consequential movements of the vocal tract, we append complementary speech features that improve reconstruction of original speech. In addition to voicing and intensity of the speech signal, we added place manner tuples (represented as continuous binary valued features) to bootstrap the EMA with what we determined were missing physiological aspects in EMA. There were 18 additional values to capture the following place-manner tuples: 1) velar stop, 2) velar nasal, 3) palatal approximant, 4) palatal fricative, 5) palatal affricate, 6) labial stop, 7) labial approximant, 8) labial nasal, 9) glottal fricative, 10) dental fricative, 11) labiodental fricative, 12) alveolar stop, 13) alveolar approximant, 14) alveolar nasal, 15) alveolar lateral, 16) alveolar fricative, 17) unconstructed, 18) voicing. For this purpose, we used an existing annotated speech database (Wall Street Journal Corpus) ^49^ and trained speaker independent deep recurrent network regression models to predict these place-manner vectors only from the acoustics, represented as 25-dimensional Mel Frequency Cepstral Coefficients (MFCCs). The phonetic labels were used to determine the ground truth values for these labels (e.g., the dimension “labial stop” would be 1 for all frames of speech that belong to the phonemes /p/, /b/ and so forth). However, with a regression output layer, predicted values were not constrained to the binary nature of the input features. In all, these 32 combined feature vectors form the initial articulatory feature estimates.

Finally, to ensure that the combined 32 dimensional representation has the potential to reliably reconstruct speech, we designed an autoencoder to optimize these values. Specifically, a recurrent neural network encoder is trained to convert phonological and acoustic features to the initialized 32 articulatory representations and then a decoder converts the articulatory representation back to the acoustics. The stacked network is re-trained optimizing the joint loss on acoustic and EMA parameters. After convergence, the encoder is used to estimate the final articulatory kinematic features that act as the intermediate to decode acoustics from ECoG.

### Neural decoder

The decoder maps ECoG recordings to MFCCs via a two stage process by learning intermediate mappings between ECoG recordings and articulatory kinematic features, and between articulatory kinematic features and acoustic features. We implemented this model using TensorFlow in python^50^. In the first stage, a stacked 3- layer bLSTM^16^ learns the mapping between 300 ms windows of high-gamma and LFP signals and the corresponding single time point of the 32 articulatory features. In the second stage, an additional stacked 3-layer learns the mapping between the output of the first stage (decoded articulatory features) and 32 acoustic parameters for full sentences sequences. These parameters are are 25 dimensional MFCCs, 5 sub-band voicing strengths for glottal excitation modelling, log(F0), voicing. At each stage, the model is trained to with a learning rate of 0.001 to minimize mean-squared error of the target. Dropout rate is set to 50% to suppress overfitting tendencies of the model. We use a bLSTM because of their ability to retain temporally distant dependencies when decoding a sequence^51^.

### Speech synthesis from acoustic features

We used an implementation of the Mel-log spectral approximation algorithm with mixed excitation^22^ to generate the speech waveforms from estimates of the MFCCs from the neural decoder.

### Model training procedure

As described, simultaneous recordings of ECoG and speech are collected in short blocks of approximately 5 minutes. To partition the data for model development, we allocated 2-3 blocks for model testing, 1 block for model optimization, and the remaining blocks for model training. The test sentences for P1 and P2 each spanned 2 recording blocks and comprised 100 sentences read aloud. The test sentences for P3 were different because the speech comprised 100 sentences over three blocks of freely and spontaneously speech describing picture scenes.

For shuffling the data to test for significance, we shuffled the order of the electrodes that were fed into the decoder. This method of shuffling preserved the temporal structure of the neural activity.

### Mel-Cepstral Distortion (MCD)

To examine the quality of synthesized speech, we calculated the Mel-Cepstral Distortion (MCD) of the synthesized speech when compared the original ground-truth audio. MCD is an objective measure of error determined from MFCCs and is correlated to subjective perceptual judgements of acoustic quality^22^. For reference acoustic features *mc*^(*y*)^ and decoded features *mc*^(*y*)^,

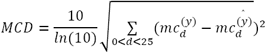

### Intelligibility Assessment

Listening tests using crowdsourcing are a standard way of evaluating the perceptual quality of synthetic speech^52^. We used the Amazon Mechanical Turk to assess the intelligibility of the neurally synthesized speech samples. We set up a listening task where naïve listeners identified which of 10 sentences was played in each trial. A set of 60 sentences (6 trials of 10 unique sentences) were evaluated in this assessment. These trials, also held out during training the decoder, were used in place of the 100 unique sentences tested throughout the rest of Figure 2 because the listeners always had the same 10 sentences to chose from. Each trial sentence was listened to by 50 different listeners. In all, 166 unique listeners took part in the evaluations.

### Data limitation analysis

To assess the amount of training data affects decoder performance, we partitioned the data by recording blocks and trained a separate model for an allotted number of blocks. In total, 8 models were trained, each with one of the following block allotments: [1, 2, 5, 10, 15, 20, 25, 28]. Each block comprised an average of 50 sentences recorded in one continuous session.

### Quantification of silent speech synthesis

By definition, there was no acoustic signal to compare the decoded silent speech. In order to assess decoding performance, we evaluated decoded silent speech in regards to the audible speech of the same sentence uttered immediately prior to the silent trial. We did so by dynamically time warping^53^ the decoded silent speech MFCCs to the MFCCs of the audible condition and computing Pearson’s correlation coefficient and Mel-cepstral distortion.

### Phoneme acoustic similarity analysis

We compared the acoustic properties of decoded phonemes to ground-truth to better understand the performance of our decoder. To do this, we sliced all time points for which a given phoneme was being uttered and used the corresponding time slices to estimate its distribution of spectral properties. With principal components analysis (PCA), the 32 spectral features were projected onto the first 4 principal components before fitting the gaussian kernel density estimate (KDE) model. This process was repeated so that each phoneme had two KDEs representing either its decoded and or ground-truth spectral properties. Using Kullback-Leibler divergence (KL divergence), we compared each decoded phoneme KDE to every ground-truth phoneme KDE, creating an analog to a confusion matrix used in discrete classification decoders. KL divergence provides a metric of how similar two distributions are to one another by calculating how much information is lost when we approximate one distribution with another. Lastly, we used Ward’s method for agglomerative hierarchical clustering to organize the phoneme similarity matrix.

## Extended Data

**Extended Data Figure 1:**
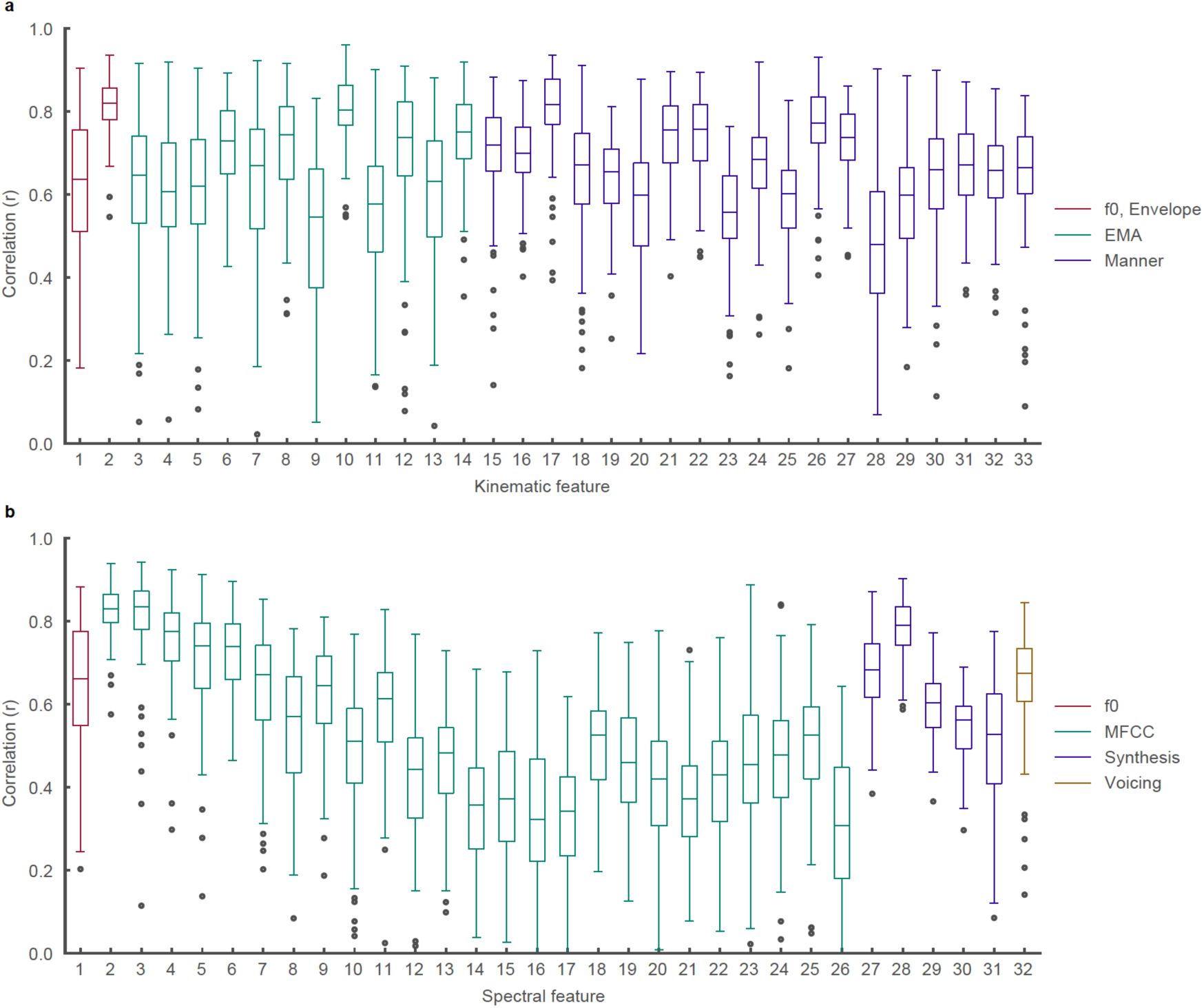
Decoding performance of kinematic and spectral features. **a**, Correlations of all 33 decoded articulatory kinematic features with ground-truth. EMA features represent X and Y coordinate traces of articulators (lips, jaw, and three points of the tongue) along the midsagittal plane of the vocal tract. Manner features represent complementary kinematic features to EMA that further describe acoustically consequential movements. **b**, Correlations of all 32 decoded spectral features with ground-truth. MFCC features are 25 mel-frequency cepstral coefficients that describe power in perceptually relevant frequency bands. Synthesis features describe glottal excitation weights necessary for speech synthesis.

**Extended Data Figure 2:**
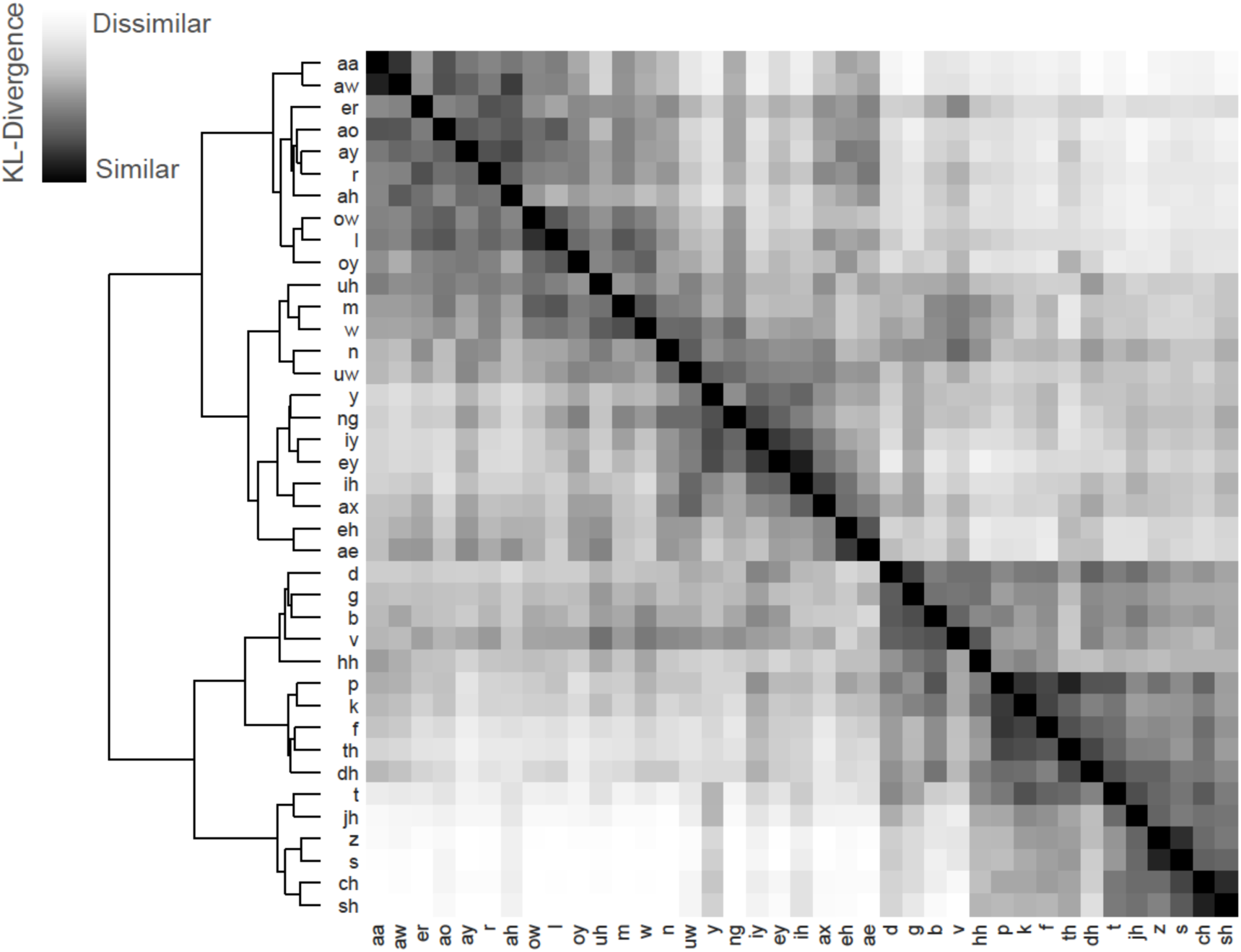
Ground-truth acoustic similarity matrix. Compares acoustic properties of ground-truth spoken phonemes with one another. Similarity is computed by first estimating a gaussian kernel density for each phoneme and then computing the Kullback-Leibler (KL) divergence between a pair of a phoneme distributions. Each row compares the acoustic properties of a two ground-truth spoken phonemes. Hierarchical clustering was performed on the resulting similarity matrix.

## Acknowledgements

We thank Matthew Leonard, Neal Fox for their helpful comments on the manuscript. We also thank Ben Speidel for his work reconstructing MRI images of patients’ brains. This work was supported by grants from the NIH (DP2 OD008627 and U01 NS098971-01). E.F.C is a New York Stem Cell Foundation- Robertson Investigator. This research was also supported by The New York Stem Cell Foundation, the Howard Hughes Medical Institute, The McKnight Foundation, The Shurl and Kay Curci Foundation, and The William K. Bowes Foundation.

## Author Contributions

Conception G.K.A., J.C., and E.F.C.; Articulatory kinematics inference G.K.A; Decoder design G.K.A and J.C.; Decoder analyses: J.C.; Data collection G.K.A., E.F.C., and J.C.; Prepared manuscript all; Project Supervision E.F.C.

## Notes

The authors declare no competing interests.

